# High-resolution electrochemistry of the extracellular electron transfer of *Escherichia coli*

**DOI:** 10.1101/2020.11.05.369678

**Authors:** Yong Xiao, Zhiyong Zheng, Haiyin Gang, Jens Ulstrup, Feng Zhao, Jingdong Zhang

## Abstract

*Escherichia coli* is one of the most important model bacteria in microorganism research and is broadly encountered in nature. In the present study, a wild-type *E. coli* strain K-12 was used for electrochemical investigations. Differential pulse voltammetry showed five pairs of redox peaks both for K-12 cells and the supernatant with potentials (anodic/cathodic) at −0.450/−0.378, −0.125/−0.105, −0.075/−0.055, +0.192/+0.264, and +0.300/+0.414 V (vs. Ag/AgCl), respectively. Chronoamperometry indicates that K-12 cells can produce immediate current by addition of glucose. The current production from K-12 can be 8-fold enhanced by 10.0 μM exogenetic vitamin K3, but addition of 10.0 μM riboflavin did not enhance the current production. Medium replacement experiments show that 50 % of the K-12 biofilm current was produced via direct extracellular electron transfer pathways. The study provides new insight in the voltammetry of strain K-12 and confirms that *E. coli* is an electrochemically active bacterium. *E. coli* has the potential to serve as a model bacterium for studying microbial extracellular electron transfer mechanisms.

## Introduction

Bioelectrochemical systems combine biological and electrochemical processes to generate electricity, hydrogen or other useful chemicals from organic materials e.g. pollutants ^1^. Microbial (EET), in which microorganisms exchange electrons with extracellular electron donors or acceptors, is here the central core process ^2-4^. A full understanding of EET mechanisms is therefore essential to improve the production of chemicals and energy as well as enhance bioremediation efficiency. Microbial EET mechanisms proposed are mainly based on studies of *Shewanella oneidensis* or *Geobacter sulfurreducens*. Redox proteins such as *c*-type cytochromes and other molecular electron shuttles, conductive pili/nanowires and extracellular polymeric substances are regarded as the core elements in extracellular transport of electrons ^5, 6^. However, it is not fully clear, how microorganisms conduct the EET process, due to the limitation of the model strains presently used: (i) *G. sulfurreducens* is an obligately anaerobic bacterium but not easy to culture, with few tools available for genetic manipulation; (ii) *S. oneidensis* is easy to culture, but genetic tools to manipulate the strain for in-depth investigations are lacking; (iii) neither of the bacteria dominate the natural environment, even in bioelectrochemical systems ^7^.

*Escherichia coli*, a facultative bacterium widely present in nature, is the most important model bacterium in microbiology and molecular biology research, with clear metabolism mechanisms mapped out and many available tools for genetic manipulation. However, there are many contradictory reports on EET ability of *E. coli*. Schröder *et al.* reported that *E. coli* K-12 produced a current of 19.5 mA through metabolic H_2_ conversion, but the electrons involved are not directly from the cells and the modified anode played an important role in the bioelectricity production ^8^. Zhang *et al.* recorded an enhanced current production from electrochemically-evolved *E. coli* K-12 together with an optimized carbon/PTFE composite anode ^9^, but did not provide a pathway for electron transfer. *E. coli* cells evolved under electrochemical tension showed electrochemical behavior in a microbial fuel cell with hydroquinone assumed to be the electron mediator ^10^. However, whether the *E. coli* K-12 of wild type have the EET ability is still widely argued; some researchers claimed that *E. coli* is a non-exoelectrogenic bacterium since the cells could not produce current in their MFCs ^11, 12^; a small residual current was ascribed to electrons and protons stored in the *E. coli* culture in a redox battery-like manner ^11^.

*E. coli* is widely abundant in soil, and understanding of *E. coli* EET mechanisms, if *E. coli* can be confirmed as an electrochemically active bacterium, can help us to illustrate and map EET processes and electrochemically active microorganisms broadly ^13^. We noted that the previous studies usually rested on biofilms grown on electrodes with large surface area and high background current. Some weak but important electrochemical signals may therefore have been missed. Hence, we have undertaken the present study of the electrochemistry of *E. coli* using a high-resolution electrochemistry method, differential pulse voltammetry (DPV) and the wild-type model strain K-12 with the view of obtaining more details on the electrochemical profile and EET ability of *E. coli*.

## Experimental Section

### Bacterial strains and growth conditions

*E. coli* K-12 strain was bought from the German Collection of Microorganisms and Cell Cultures (DSMZ) (DSM number 498). M9 medium was used to culture *E. coli* K-12 cells. The M9 medium ^14^ contains (g/L) glucose of 4.00, Na_2_HPO_4_ of 6.78, KH_2_PO_4_ of 3.00, NaCl of 5.00, NH_4_Cl of 1.00, MgSO_4_•7H_2_O of 0.493, and CaCl_2_ of 0.011. The initial pH of the M9 medium was adjusted to 7.00 using 1% HCl or 1 M NaOH. The strain was grown aerobically at 37 °C by shaking at 120 rpm. Glucose, Na_2_HPO_4_, KH_2_PO_4_, NaCl, NH_4_Cl, MgSO_4_•7H_2_O, and CaCl_2_ were analytically pure products from Sigma-Aldrich. Riboflavin and vitamin K3 were USP Grade from Sigma-Aldrich and from Coolaber Inc., China, respectively. *E. coli* K-12 cells were harvested by centrifugation (5000 *g*, 10 min, 4 °C) and washed twice with 0.05 M phosphate potassium buffer (pH 7.00) before use.

The *E. coli* K-12 strain grew fast aerobically in M9 medium by shaking at 125 rpm and reached the stationary phase in 12 h (Figure S1.A). Cells grown at 12-16 h with a length of around 2 μm (Figure S1.B and C) were harvested from the M9 medium for voltammetric recording by DPV, CV, and chronoamperometry.

### Electrochemical measurements on *E. coli* K-12 cells

All electrochemical voltammetry experiments were carried out using a CHI832D or CHI732D potentiostat (CHI, TX, USA) with a three-electrode chamber containing a glassy-carbon working electrode (3 mm diameter), a platinum-wire counter electrode and a KCl saturated Ag/AgCl reference electrode ^6, 15^. All potentials reported refer to Ag/AgCl sat. KCl. Phosphate potassium buffer solution of 0.05 M (pH 7.00) was used as electrolyte. All experiments were conducted under a nitrogen atmosphere. The parameters were: for cyclic voltammetry (CV), equilibrium time 5 s, scan rate 10 mV/s; for differential pulse voltammetry (DPV), equilibrium time 5 s, increment potential 6 mV, amplitude 60 mV, pulse width 0.2 s, pulse period 0.4 s; for *i-t* chronoamperometry, the working electrode was kept at +0.10, +0.20, or +0.30 V as indicated in each figure legend.

### *E. coli* K-12 biofilm

*E. coli* K-12 biofilm was anaerobically grown on carbon felt (area 1.0 cm^2^) which also served as the working electrode for subsequent *i-t* chronoamperometry. Stainless steel mesh (area 9.0 cm^2^) and a KCl saturated Ag/AgCl electrode served as counter and reference electrode, respectively. The electrodes were connected to a CHI1000C potentiostat (CHI, TX, USA). The electrode system was inserted in a sealed 100 mL flask filled with oxygen-free medium during the cultivation. A polypropylene tube was connected to the bottom of the flask when the medium was replaced. Magnetic stirring (50 rpm) was also used.

## Results and Discussion

### Voltammetry and chronoamperometry of *E. coli* K-12

Five pairs of DPV redox peaks of *E. coli* K-12 cells grown at 12-16 h i.e. the early stationary growth phase (Figure 1A) were observed. The potentials of the anodic peaks were −0.450, −0.125, −0.075, +0.192, and +0.300 V. The corresponding cathodic peak potentials were −0.378, −0.105, −0.055, +0.264, and +0.414 V. The redox peaks around −0.400 V, −0.200 to 0 and 0.200 to 0.400 V were grouped into the redox peak group I, II and III, respectively (Figure 1A). The potentials of the group II and III redox peaks were so close that they sometimes overlap into a single redox peak (Figure S2). Only one or two anodic peaks were observed in the voltammetry of biofilms on electrodes with large surface area, for example −0.100, +0.100 V ^16^ or −0.070 V ^17^. By conducting voltammetry on collected *E. coli* K-12 cells on a glassy carbon electrode, the data in the present study displayed, however, a notably more sensitive *E. coli* electrochemical redox profile than in previous reports ^16, 17^.

**Figure 1.**
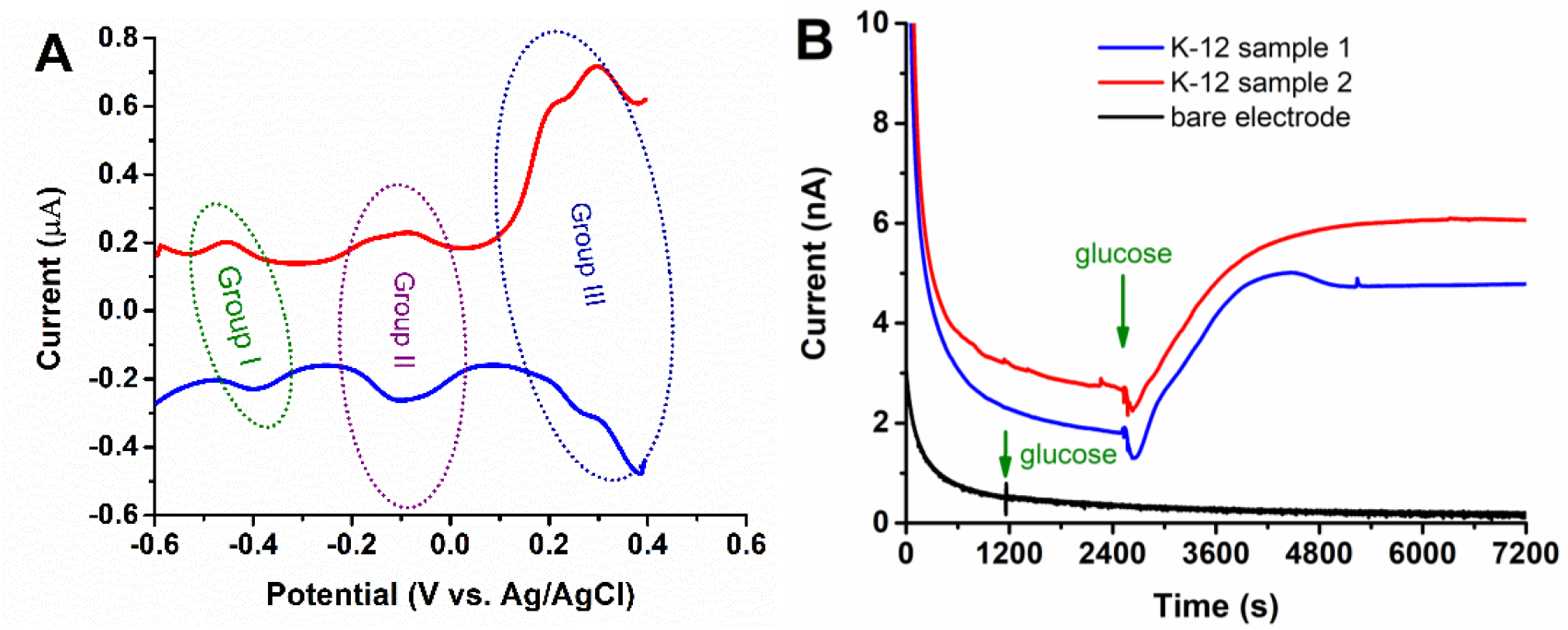
Electrochemical patterns of *E. coli* K-12 cells grown at 16 h. (A) DPV of *E. coli* K-12 cells dropcast onto a glassy carbon electrode. Five redox peak pairs were found and grouped into three groups. While there is only one redox peak pair in group I, group II and III each contains two pairs of redox peaks. (B) Chronoamperometry confirms a current response to added glucose from *E. coli* K-12 cells when the working electrode was polarized at +0.10 V. 3.00 mL 40.0 g/L glucose solution was added to 27.00 mL electrolyte, making the final concentration 4.00 g/L glucose. No current was produced from bare glassy carbon electrode in 4.00 g/L glucose. Glassy carbon, Ag/AgCl, and platinum wire served as working, reference and counter electrode, respectively. In 0.05 M phosphate potassium buffer of pH 7.00.

All the five redox peak pairs observed for *E. coli* K-12 cells were also found in the CV profiles of the supernatant (Figure S3). As the scans continued, the group II current increased, whereas those in group III decreased. No obvious change was observed in group I. These results indicate that the electroactive molecules on the cell membrane are also excreted to the supernatant. Furthermore, it seems that the molecules related to group II are easily adsorbed on glassy carbon while those related to group III are very unstable, since the peak current of this group continued to decrease.

Chronoamperometry confirmed that *E. coli* K-12 cells can produce current and have EET ability using glucose as electron donor at a polarization potential of +0.10 V, whereas glucose cannot be used for current production without cells (Figure 1B). Compared to glucose, *E. coli* K-12 cells used lactose for a smaller current production while they did not produce current from lactate and acetate 1 h following substrate addition (Figure S4). Furthermore, biofilm formed anaerobically on carbon felt (1.0 cm^2^) can produce a current increment of 0.20 μA by supplying glucose at a polarization potential of +0.20 V (Figure S5). These results confirm that *E. coli* is an electrochemically active bacterium, able to produce electricity from appropriate organic substrate such as glucose, and redox peak groups I and II may both contribute to the electricity production based on the hold potential of +0.20 V.

### What is the role of riboflavin in EET?

The pair of redox peaks around −0.40 V i.e. group I is ascribed to flavins as the redox potentials are similar to flavins potentials in many other microorganisms such as *Shewanella* spp. ^6, 18, 19^, *Bacillus* spp. ^20, 21^, *Listeria monocytogenes* ^22^, *Pichia stipitis* ^20^, and *Pachysolen tannophilus* ^23^. Previous studies also reported that *E. coli* can excrete flavins during growth with glucose ^24^. Besides, we found that the redox peaks are strengthened by riboflavin addition (Figure S6), meaning that the two substances have similar redox potentials.

Flavins have been widely reported as ET mediators for microbial EET process ^6, 20, 21, 23^, but low concentration of riboflavin cannot be used by *E. coli* K-12 cells to facilitate its EET as no current increment was observed after the addition of extra 10.0 μM riboflavin (Figure 2). This result agrees well with a previous study which showed that *E. coli* secreted three types of flavins ^19^, but that *E. coli* did not utilize these molecules in EET ^25^. The result further confirmed that the electric current produced by *E. coli* K-12 at polarization potentials +0.10 or +0.20 V are mainly contributed by the redox peak group II, but not by group I ascribing to flavins.

**Figure 2.**
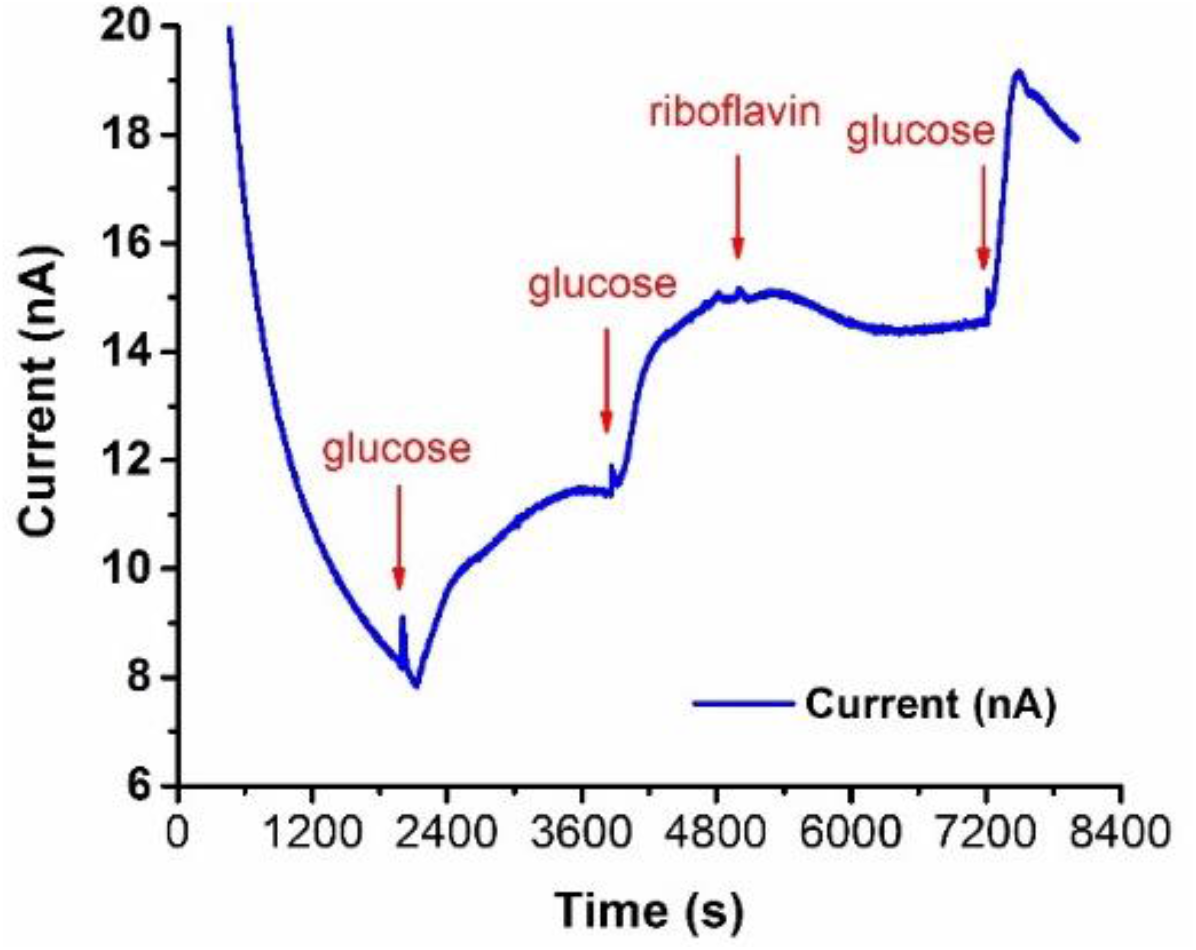
Chronoamperometry of *E. coli* K-12 cells dropcast on glassy carbon electrode. This result shows a fast current response to each glucose addition from *E. coli* K-12 cells when the working electrode was polarized at +0.20 V. Each time glucose with a final concentration of 1.00 g/L increase was added. However, no current increment was observed after the addition of riboflavin (final 1.0 μM). The Ag/AgCl and platinum wire served as reference and counter electrode, respectively. 0.05 M phosphate buffer of pH 7.0.

### Effect of vitamin K3 on the *E. coli* EET

Previous reports suggested that *E. coli* may use menaquinone or related compounds as mediators for EET ^17^, and the bacterium can definitely produce vitamin K2, i.e. menaquinone ^26^. However, vitamin K2 is not soluble in water and it is not obvious, how *E. coli* can utilize this substance as an electron shuttle in solution. One possible explanation if the compound is truly a K2 molecule, is that this molecule may combine with another molecule or even proteins. In the present study, we added another soluble synthetic vitamin K3 (final concentration 10.0 μM) with the same functional group *i.e.* the quinone group to test whether *E. coli* K-12 cells can use it as a mediator during the CV of *E. coli* K-12. The result shows that compared with the redox peak group II, vitamin K3 gave more negative redox peak potentials and definitely strengthened the oxidative current but weakened the reduction current of the *E. coli* K-12 strain (Figure 3A).

**Figure 3.**
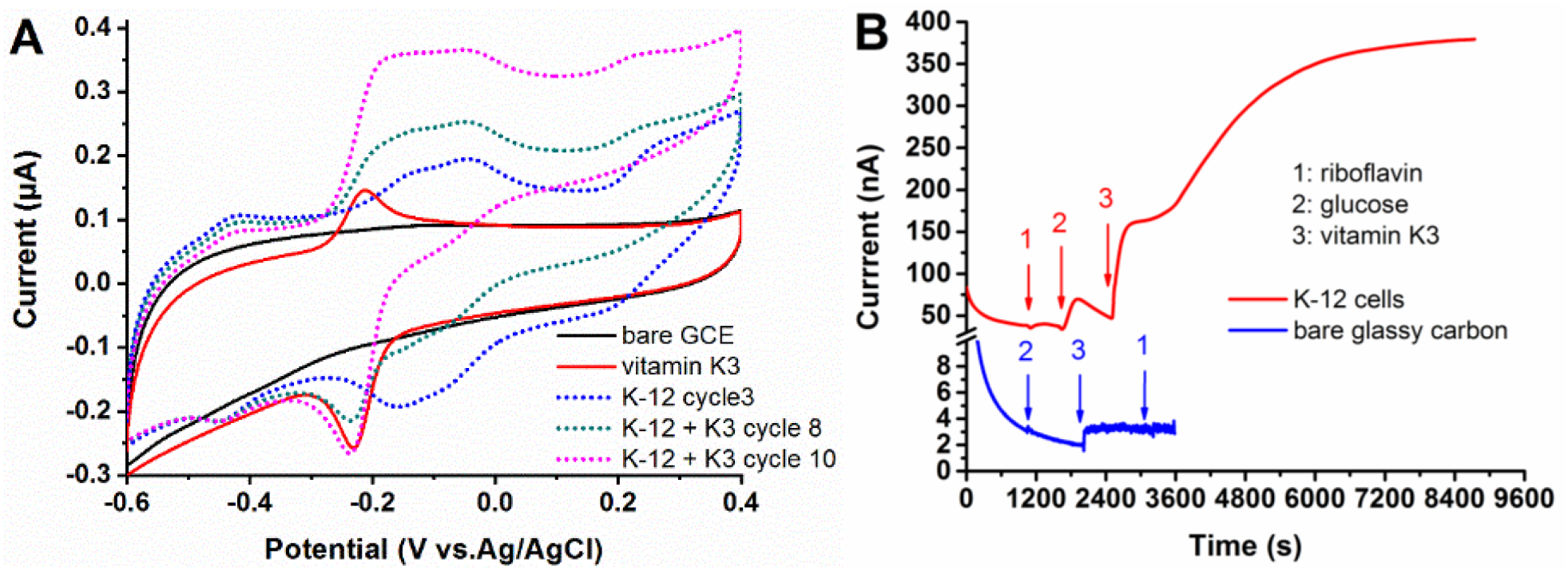
Effects of vitamin K3 on the EET of *E. coli* K-12. (A) CV of vitamin K3 and *E. coli* K-12 cells grown for 16 h with and without extra vitamin K3 added. 10.0 μM vitamin K3 showed a pair of redox peaks around −0.200 V (red line). K3 (final 10.0 μM) was added to the electrolyte at the beginning of cycle 7. The original CV pattern (blue dotted line, data from cycle 3) showed the three groups of redox peaks from *E. coli* K-12. The presence of K3 gradually strengthened the oxidative current beginning at −0.250 V (green dotted line and purple dashed line, data from cycle 8 and 10, respectively) as it diffused into the biofilm on the glassy carbon surface. Scan rate 10 mV/s. (B) Chronoamperometry shows that *E. coli* K-12 cells can use vitamin K3 but not riboflavin to enhance EET. Only a very small current increment from bare glassy carbon on addition of vitamin K3 was observed. Glassy carbon, Ag/AgCl and platinum wire served as working, reference and counter electrodes, respectively. 0.05 M potassium phosphate buffer, pH 7.00. The working electrode potential was kept at +0.30 V. The final concentration of riboflavin, glucose and vitamin K3 were 10.0 μM, 4.00 g/L and 10.0 μM, respectively.

We further evaluated the impact of vitamin K3 on the current production of *E. coli* K-12 from glucose and compared with that of riboflavin by sequentially adding riboflavin, glucose and K3 into the electrolyte (Figure 3B). Riboflavin apparently had very little positive effect on the EET of *E. coli* K-12 cells, and we only observed a very small current increment (from 35.5 to 40.1 nA) caused by the addition of 10.0 μM riboflavin. The current increased (from 33.8 nA to a maximum of 69.8 nA) by sequential addition of 4.00 g/L glucose, which agreed well with the result in Figure 1B. As a contrast, vitamin K3 was added into the electrolyte when the current was decreasing. The current continued to increase very fast from 46.8 nA to more than 380.0 nA. A current increment of only about 1.0 nA was observed when the same amount of glucose and vitamin K3 was added to a similar system but without *E. coli* K-12 cells on the working electrode (Figure 3B).

Vitamin K3 can also boost current from *E. coli* K-12 biofilm formed on carbon felt (area about 1.0 cm^2^, Figure S7). These results indicate that *E. coli* K-12 cells can use water soluble vitamin K3 as a mediator or combine this molecule with something on the cell surface to promote significantly the EET process and current production. However, whether *E. coli* can use vitamin K2 with the same quinone group as that in vitamin K3 to facilitate the EET process is not clear.

### Direct or indirect mechanisms dominate the EET process of *E. coli* K-12 cells?

Following a previous study on *S. oneidensis* MR-1 ^18^, in the present study, we have employed medium replacement to identify which, direct or indirect pathway, dominates the EET process of *E. coli* K-12 cells. As shown in Figure S8, about 50% oxidative current remains after the original supernatant is replaced by oxygen-free fresh M9 medium. These results were entirely reproducible both in 60 and 18 hold *E. coli* K-12 biofilms formed on carbon felt. The results indicate that direct and indirect electron transfer mechanisms are of equal importance in the EET of *E. coli* cells. Since flavins do not contribute to current production, the direct as well as the indirect electron transfer,of *E. coli* should be contributed by the redox peak groups II and III. However, more in-depth work including molecular biology and electrochemistry are needed to specify the contribution of each redox peak to direct or indirect electron transfer.

## Conclusions

By using small working electrodes and highly sensitive DPV measurements, high-resolution voltammetric profiles of *E. coli* K-12 were acquired in the present study. Five pairs of DPV redox peaks in the electrochemical window −0.600 to +0.400 V were disclosed. The redox peaks around −0.400 V (the group I peaks) belong to flavins which could not be utilized by *E. coli* K-12 cells to facilitate their EET processes. *E. coli* K-12 cells can instead use water soluble vitamin K3 to enhance the oxidative current from glucose. Direct and indirect electron transfer mechanisms are of equal importance in the EET of *E. coli* K-12 cells. Though wildtype *E. coli* K-12 produces relative small currents, this study confirms *E. coli* as an electrochemically active bacterium. To fully illustrate the EET process of *E. coli*, future combined work from electrochemistry and molecular biology, can be performed to elucidate the origins of the other four pairs of redox peaks. Such information would also boost the application of *E. coli* in bioelectrochemical systems and bioelectricity production.

## Acknowledgements

Financial support from the National Natural Science Foundation of China (51878640, 51478451), the Youth Innovation Promotion Association of Chinese Academy of Sciences (2018344), the Carlsberg Foundation (CF15-0164), China Scholarship Council ([2016]3100), Universities Denmark and Otto Mønsted Foundation is gratefully appreciated.

## Contributions

Y.X., F.Z. and J.Z. designed the experiments. Y.X., Z.Z. and H.G. performed the experiments. Y.X. drafted the manuscript. All the authors revised and approved the manuscript.

