## supplemental information for "High-resolution electrochemistry of the extracellular electron transfer of *Escherichia coli*"

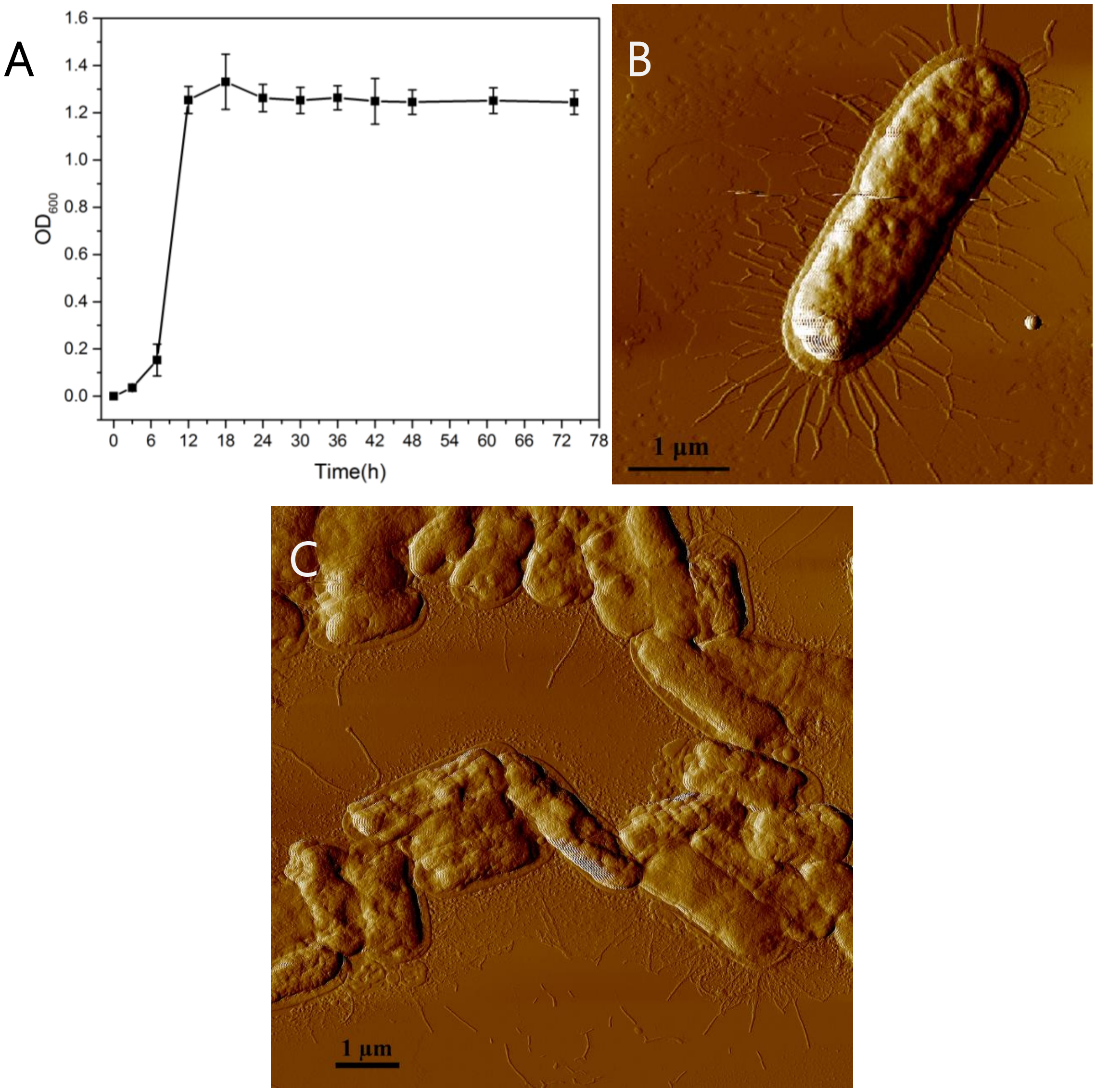


**Figure S1 (A) Growth curve of *E. coli* K-12 in M9 medium at 37 °C by shaking at 120 rpm; (B) and (C) the morphology of *E. coli* K-12 cells on mica mapped by atomic force microscopy (AFM 5500, Agilent, USA).** Individual *E. coli* cells with many micrometer scale pili are clearly visible.

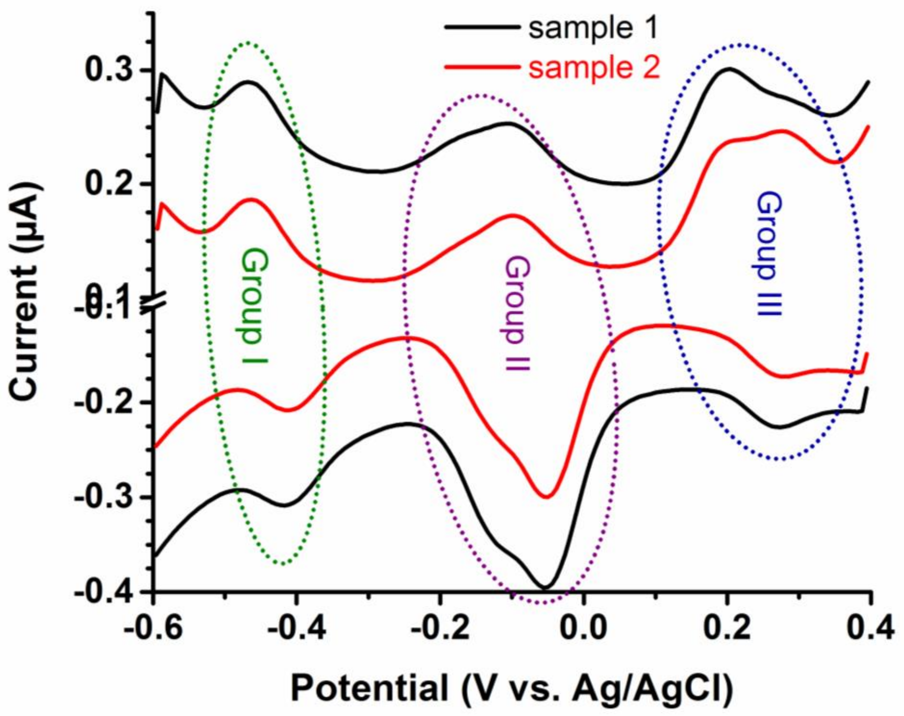


**Figure S2** **DPV profiles of K-12 cells after 12 h growth.** The oxidative peaks in group II from sample 2 and III from sample 1 are not well separated. For DPV, equilibrium time 5 s, increment potential 6 mV, amplitude 60 mV, pulse width 0.2 s, pulse period 0.4 s. Phosphate buffer (0.05 M, pH 7.00). Glassy carbon, Ag/AgCl and platinum wire served as working, reference and counter electrodes, respectively.

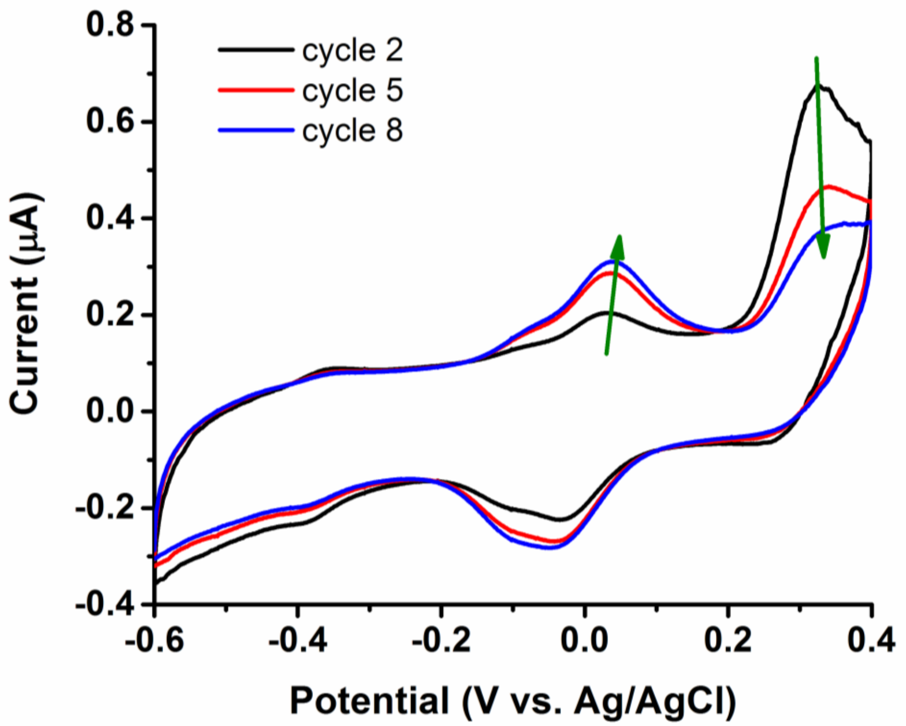


**Figure S3 CV profiles of supernatant collected from K-12 strain after 16 h cultivation at 37 °C and shaking at 120 rpm.** Bare glassy carbon, Ag/AgCl and platinum wire served as working, reference and counter electrodes, respectively. pH of supernatant 6.03. Scan cycle 2 (black), 5 (red) and 8 (blue) are shown. Scan rate 10 mV/s.

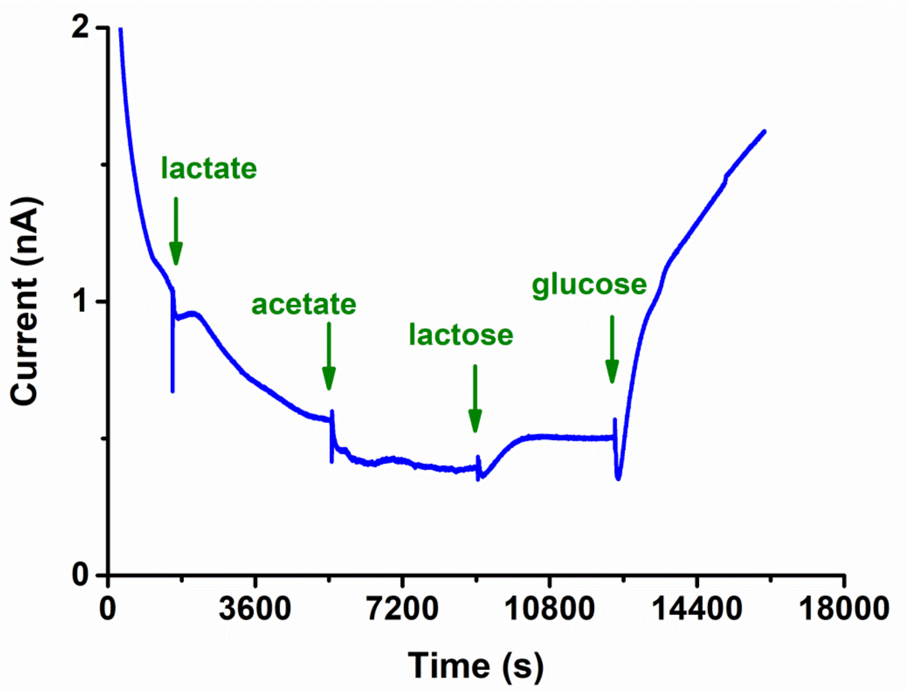


**Figure S4** ***C*****hronoamperometry of *E. coli* K-12 cells on glassy carbon electrode.** Glassy carbon, Ag/AgCl, and platinum wire served as working, reference and counter electrodes, respectively. Lactate, acetate, lactose and glucose (each final concentration 4.00 g/L) was added to the phosphate buffer (0.05 M, pH 7.00) at time points 1800, 5400, 9000, and 12600 s. The glassy carbon working electrode was polarized at +0.10 V (vs. Ag/AgCl).

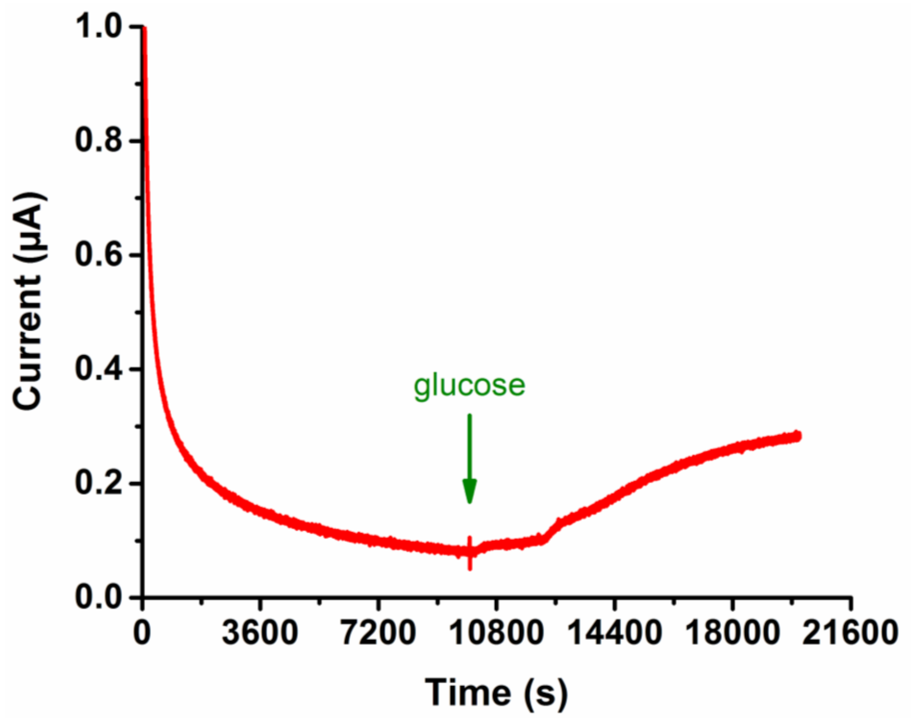


**Figure S5** ***i-t* chronoamperometry on biofilm formed on carbon felt.** The biofilm was grown on carbon felt (1.0 $cm^2$) in M9 medium for 60 h and then transferred to 0.05 M phosphate buffer (pH 7.00). The biofilm served as working electrode. An Ag/AgCl electrode and stainless steel mesh (9.0 $cm^2$) served as reference and counter electrode, respectively. The final concentration of glucose was 4.00 g/L. The potential was kept at +0.20 V.

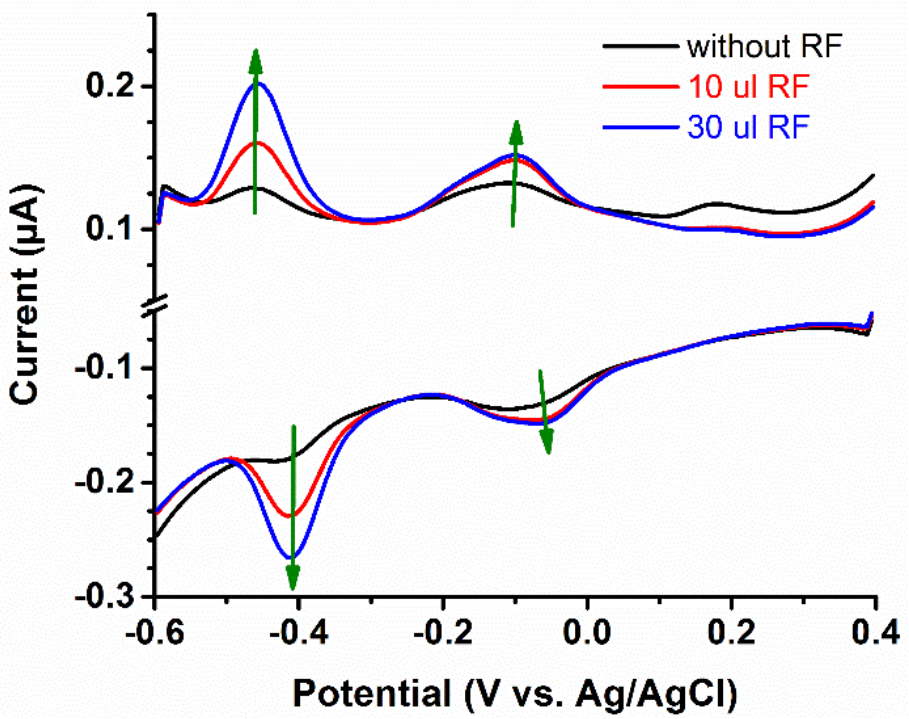


**Figure S6 DPV profiles of K-12 cells grown at 24 h without or with exogenetic riboflavin (RF) added.** Glassy carbon, Ag/AgCl, and platinum wire served as working, reference and counter electrodes, respectively. The profiles show that the current of the redox peak at potential about -0.40 V is increased by adding 10.0 (red line) and another 20.0 (blue line) µl 3.0 mM RF into 30.00 mL 0.05 M phosphate buffer (pH 7.00), i.e. the final RF concentration is 1.0 µM.

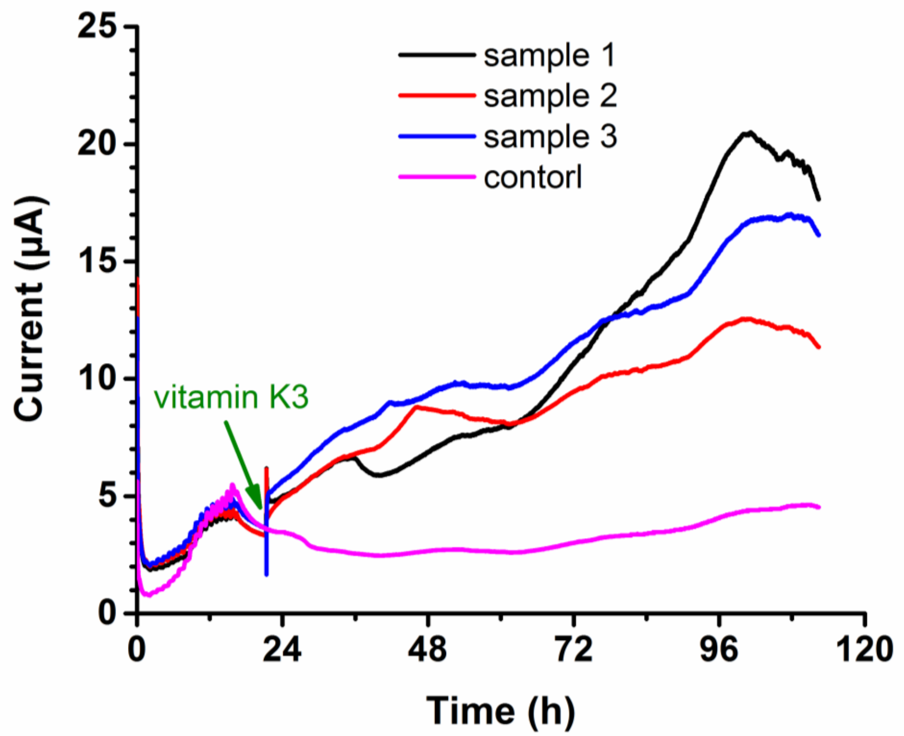


**Figure S7** ***i-t* chronoamperometry showing that the current from the biofilm formed on carbon felt could be boosted by adding 10.0 μM vitamin K3.** "Vitamin K3" indicates the addition of vitamin K3 in sample 1, 2, and 3. No vitamin K3 was added in the control sample. The area of the carbon felt i.e. the working electrode was 1.0 $cm^2$. A Ag/AgCl electrode and stainless steel mesh served as reference and counter electrode, respectively. The polarization potential was +0.20 V. The medium was M9 medium.

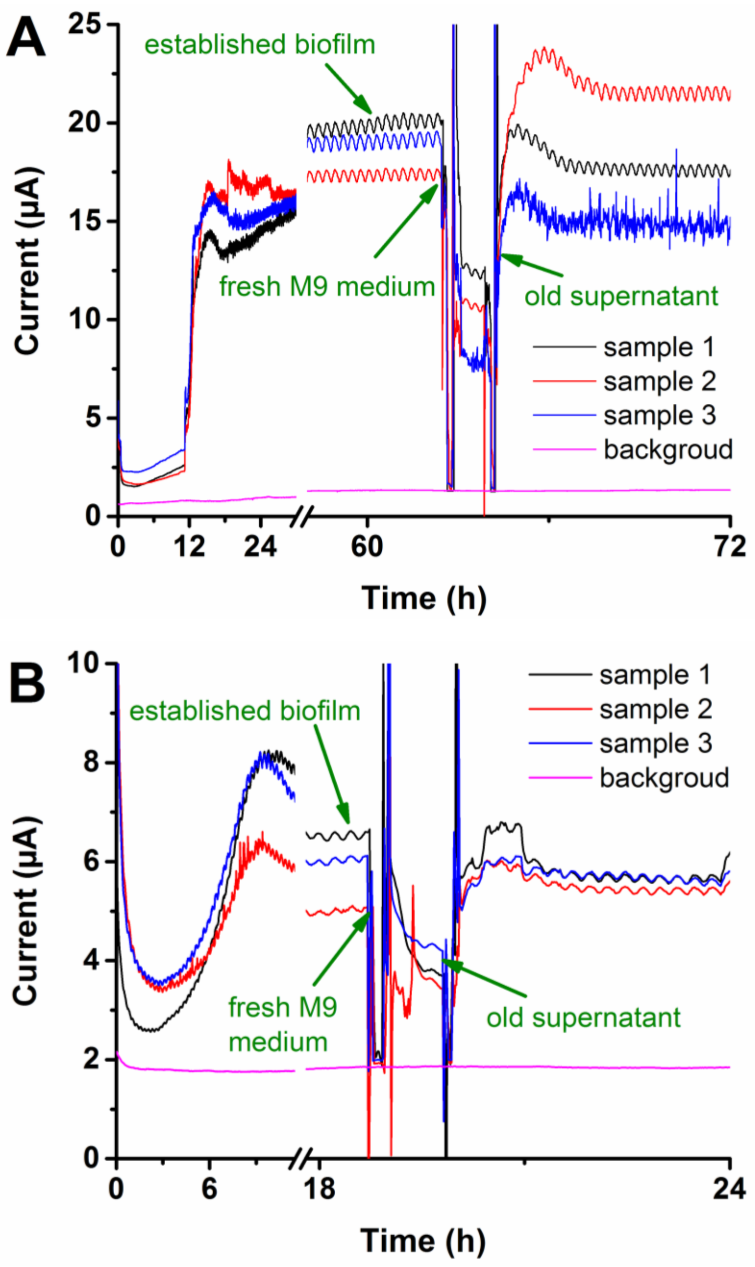


**Figure S8 Medium replacement showing that about 50% of the current remained after the supernatant has been removed and fresh M9 medium pumped in.** (A) Medium replacement on 60 h-old K-12 biofilm. (B) Medium replacement experiment on 18 h-old K-12 biofilm. The area of the working electrode in both A and B was about 1.0 cm$^2$. A Ag/AgCl electrode and a stainless steel mesh served as reference and counter electrode, respectively. The potential was kept at +0.30 V. The medium was M9 medium. Oxygen was removed from the fresh M9 medium by bubbling $N_2$ for 30 min. The ‘established biofilm’ represents a biofilm formed on carbon felt. The ‘fresh M9 medium’ represented the process of removing old supernatant and pumping in fresh M9 medium, and the ‘old supernatant’ represents the process of removing fresh M9 medium and pumping in old supernatant. The ‘background’ current was obtained from a similar three-electrode system without K-12 cells. The waves on the current curves are induced by the temperature fluctuations in the water bath.
